# VIRAL METAGENOMIC SURVEY OF CASPIAN SEALS

**DOI:** 10.1101/2024.07.14.603418

**Authors:** K Karamendin, SJ Goodman, Y Kasymbekov, M Kumar, Nuralibekov, A Kydyrmanov

**Affiliations:** Laboratory of Viral Ecology, Scientific and Production Center of Microbiology and Virology, Department of Virology, Almaty, Kazakhstan; School of Biology, Faculty of Biological Sciences, University of Leeds, Leeds, United Kingdom

**Keywords:** Caspian seal, Pusa capsica, viral metagenome, marine mammals, pinniped, virome, wildlife diseases

## Abstract

Viral diseases of pinnipeds cause substantial mortality and morbidity and can influence population demography. Viral metagenomic studies can therefore play an important role in pinniped health assessments and disease surveillance relevant to both individual species and in a ‘One Health’ context. This study used a metagenomic approach with high throughput sequencing to make the first assessment of viral diversity in Caspian seals (*Pusa caspica*), the only marine mammal species endemic to the Caspian Sea. Sequencing libraries from 35 seals sampled 2009-2020 were analysed, finding sequences from the viral families *Picornaviridae, Adenoviridae, Circoviridae, Herpesviridae, Papillomaviridae, Caliciviridae, Orthomyxoviridae, Anelloviridae, Smacoviridae, Cruciviridae* and *Parvoviridae*. The similarity of viral contigs from Caspian seal to sequences to those recovered from other pinnipeds ranged from 63.74% (San Miguel sea lion calicivirus) to 78.79% (Seal anellovirus 4). Some may represent novel viral species, but overall, the viral repertoire of Caspian seals is similar to available viromes from other pinnipeds. Among the sequences recovered were partial contigs for influenza B, representing only the second such molecular identification in marine mammals. This work provides a foundation for further studies of viral communities in Caspian seals, the diversity of viromes in pinnipeds more generally, and contributes data relevant for disease risk assessments in marine mammals.

## Introduction

Marine mammals are sentinels for ocean health [1]. Monitoring the health status of wild marine mammal populations is essential for understanding the exposure and vulnerability of individual species to pathogens, the welfare of individuals and population dynamics, as well as the implications for the overall resilience of marine ecosystems to environmental change and anthropogenic threats [2, 3]. Studies of marine mammal health are also relevant to the ‘One Health’ context due to their interactions with humans and other species, and because anthropogenic activities may influence pathogen exposure risk. Long term monitoring suggests that in North America between 1991 and 2022, infectious diseases accounted for 14% of marine mammal deaths, with biotoxins (18%), ecological factors (14%), human interactions (5%), and other undetermined factors (49%) contributing the remainder [4]. Infectious disease pathogens (viruses, bacteria and parasites), can influence individual mortality and morbidity, with consequences for survival and reproductive output, and drive mass mortality events, all of which can significantly impact marine mammal population demography [5]. Therefore, it is important to develop baseline information about pathogen repertoires and prevalence to guide development of surveillance programmes, to assess temporal changes, and facilitate investigations of mass mortalities in support of marine mammal conservation and management [6].

Knowledge of the diversity of marine mammal pathogens is expanding rapidly [7-12], with morbilliviruses [5, 6] and influenza viruses [13, 14] being some of the best documented drivers of mass mortality events. The development of high throughput deep sequencing is driving new capability for pathogen discovery and surveillance beyond that achievable with standard serological and genetic approaches [15]. In the context of viruses, metagenomic virome studies sequence the total nucleic acid profile from clinical or necropsy samples to assemble and identify sequence contigs from any pathogens present. Comparing recovered sequences to genomic databases, both known and novel viruses can be identified. Viromes for some members of the *Otariidae* family *(eared seals)* have been well characterised. Studies in California sea lions (*Zalophus californianus*) found previously undescribed RNA virus families for the species, including *Astroviridae, Picornaviridae, Caliciviridae*, and *Reoviridae*, and one DNA virus family (*Parvoviridae*) [16]. An analysis of the virome of subantarctic fur seals (*Arctocephalus tropicalis*) and South American fur seals (*Arctocephalus australis*) has revealed the sequences of adenoviruses, circoviruses, picobirnaviruses, picornaviruses, herpesvirus-like viruses, anelloviruses, parvoviruses and one putative new genus, named *Nitorquevirus* [15, 17, 18]. For phocid (earless) seals, metagenomic studies are less extensive, but have contributed to the discovery of novel anelloviruses, circoviruses, cycloviruses and papillomaviruses in Weddell seals (*Leptonchotes weddelli*) [19-22]; papillomaviruses in Leopard seals (*Hydrurga leptonyx*) [21]; and anelloviruses [23], varicellovirus (PHHV-1) [24], and gammaherpesvirus (PhHV-7) [25] in harbour seals (*Phoca vitulina*).

The Caspian seal (*Pusa caspica*) is a small-bodied phocid seal endemic to the Caspian Sea [26], and is classified as ‘Endangered’ by the International Union for Conservation of Nature (IUCN) Red List, with a population size of around 168,000 [27]. Historically, mass mortalities in Caspian seals were first recorded in 1955-56, when about 30 thousand animals died from a purported bacterial infection [28]. However, at the time, the capacity to detect viral infections was limited, and the role of bacterial secondary infections under-appreciated. Further mass mortalities occurred in the Kazakh part of the Caspian Sea in 1968, 1971, 1978 and 1985, with the cause of death attributed to underwater blasting connected with hydro-engineering and military works [29], but with no toxicological, epidemiological and virological studies conducted. In 1997-2000 several large-scale seal stranding events were observed on the Caspian coast, with total mortality exceeding more than 11,000 seals [30]. Canine distemper virus, a *Morbillivirus*, was identified as a primary causative agent [31]. Since 2000, regular mass strandings have been reported ranging from 10s to 1000s of individuals, but the causes remain poorly investigated due to challenging logistics and limited capacity for marine mammal pathology in the region [32], [33].

The Caspian seal, as a transboundary species, is a key indicator of the state of the Caspian ecosystem [34]. In order to develop evidence-based conservation strategies it is important to be able to evaluate the effects of infectious diseases in the context of other anthropogenic factors such as impacts from shipping [35], fisheries bycatch [36], pollution [32], and historical hunting [37].

Kydyrmanov et al. recently reported on the long-term screening of prevalence for viruses, bacteria and parasites in Caspian seals, sampling 177 live, healthy, wild Caspian seals (Pusa caspica) between 2007 and 2017, using serological and PCR based methods, concluding the overall pathogen repertoire and prevalence was comparable to other phocid seal populations [33]. In that work, evidence for viral infections included CDV, phocine herpes virus, phocine adenovirus, influenza A, influenza B, and coronavirus. Here we report the first investigation of the virome of the Caspian seal using a metagenomic approach. The results add new knowledge on the viral epidemiology of this species, and will contribute to the understanding of current and future disease threats.

## Materials and Methods

### Seal capture and sample collection

Samples for this study were collected during fieldwork in the Kazakh part of the northeastern and central Caspian Sea (Kazakh coastline and nearby islands) during spring and autumn of 2009, 2016, 2019 and 2020 (Supplementary Table S1). The seal haul out sites were reached by rigid inflatable boats. After stealthily approaching the seals on land, animals were caught using a ‘rush and grab’ approach with hoop nets. Animals were restrained and handled without the use of chemical sedation, and were released immediately after the completion of sampling [33]. Sampling was conducted in accordance with the Rules for conducting biomedical experiments, preclinical (non-clinical) and clinical studies (№697, 12 November 2007, Republic of Kazakhstan and Local Ethics Committee Research and Production Center for Microbiology and Virology (Approval number: 02-09-05 from 19 August 2014). In December 2020, about three hundred seals were reported stranded dead along the coast of Dagestan in Russia [38]. Tissue samples (lung, spleen, kidney, large intestine) and swabs (tracheal, nasal, rectal) from eight carcasses from this event, stored in DNA/RNA shield were delivered to the laboratory and then archived at 4 °C.

Body length (from nose tip to tail tip), girth and weight, and sex was recorded for each individual. Seals were categorised as juveniles (<1 year; body length 70–90 cm), sub-adults (older than one year of age; length >91–109 cm), adults (mature, length >110–140 cm). A complete external body examination was made for skin ulcers, parasites, trauma lesions and any visible alterations. Duplicate nasal, buccal, rectal, and urogenital samples were collected from captured seals using sterile cotton swabs, following Marine Mammal Commission guidelines [39]. For virus isolation, swabs were placed in vials with viral transport medium (VTM) 199 containing antibiotics (penicillin 2000 U/ml, streptomycin 2 mg/ml, gentamicin 50 μg/ml, nystatin 50 U/ml) and bovine serum albumin at a final concentration of 0.5%. The samples were stored in liquid nitrogen (–196°C) until delivery to the laboratory. For molecular analysis, swabs were placed in RNA/DNA Shield reagent (Zymo Research, USA), which preserves viral nucleic acids at ambient temperatures.

### Sample Processing

Before nucleic acid extraction, swab samples in VTM were centrifuged at 3200 RPM for 15 minutes and the resulting liquid was passed through a 0.22 µm filter (Membrane Solutions, USA) and then the filtrate was treated with a mix of nucleases: Benzonase (Sigma-Aldrich, USA), Turbo DNAse, DNAse I, RNAse A, and RNAse T1 (ThermoFisher, Lithuania). Solid tissue samples (5mm^3^) were placed in 2-ml vial with sterile PBS, pH 7.4 (Sigma Aldrich, USA) and beads, homogenized with a TissueLyser instrument (Qiagen, Germany) shaking with a frequency of 25 cycles per second for 3 min, and centrifuged at 3200 RPM for 15 minutes to pellet the debris. Viral nucleic acids were then extracted from swab filtrates and tissue homogenate supernatants using the QIAamp Viral RNA Mini Kit (Qiagen, Germany) following the manufacturer’s recommendations. The QIASeq RNA Kit (Qiagen, Germany) was then used to generate double-stranded cDNA from input RNA.

### Sample pooling

Extractions from 35 seals were organised into 34 pools of different sample type combinations for sequencing library construction (Table 1; Supplementary Table 1). For material collected in Kazakhstan from 2016, 2019 and 2020, libraries consisted of either individual or pooled samples. Within years, extractions from the same sample type (buccal, nasal, genital, rectal and conjunctival swabs) were pooled by sex-age class (juvenile males, adult males, juvenile females, and adult females). Additionally, one library was constructed from a buccal swab from an adult female sampled in 2009, which was PCR-positive for avian influenza A [33]. Lastly, a library was produced for a necropsy spleen sample from one of the Dagestan seals from the 2020 mass mortality event.

**Table 1.** Year/Sex/Age class composition of sample pools and total contig counts for each detected virus family found in each pool.

| Sample ID | Age | Sex | Sample Type | Total Contigs of Viral Families Found |  |  |  |  |  |  |  | Ortho-myxo- | Cruci- |
| --- | --- | --- | --- | --- | --- | --- | --- | --- | --- | --- | --- | --- | --- |
|  |  |  |  | Circo- | Herpes- | Parvo- | Papillo- | Picorna- | Calici- | Anello- | Smaco- |  |  |
| Seal-Flu-2009-11* | Ad | F | B |  |  |  |  |  |  |  |  | 5 |  |
| Seal_2016-3 | Ad | M | N | 23 | 15 |  |  |  |  |  |  |  |  |
| Seal_2016-4B | Ad | F | B | 1 |  |  |  | 3 |  |  |  |  |  |
| Seal_2016-4G | Ad | F | G | 2 |  |  |  | 14 |  |  |  |  |  |
| Seal_2016-4N | Ad | F | N |  |  |  |  | 2 |  |  |  |  |  |
| Seal_2016-4R | Ad | F | R | 16 |  |  |  | 82 |  |  |  |  |  |
| Seal_2016-5G | Ad | F | G | 2 |  |  |  |  |  |  |  |  |  |
| Seal_2016-6R | Ad | F | R |  |  |  |  | 2 |  |  |  |  |  |
| Seal_2016-16R | Ad | F | R |  |  |  |  | 6 |  |  |  |  |  |
| Seal_2019-pool-1 | Ad | M | B |  | 15 |  |  |  | 11 |  |  |  |  |
| Seal_2019-pool-2 | Juv | F | B |  | 22 |  |  |  |  |  |  |  |  |
| Seal_2019-pool-3 | Ad | F | B |  | 10 |  |  |  |  |  |  |  |  |
| Seal_2019-pool-4 | Ad | M | G |  | 25 |  |  |  |  |  |  |  |  |
| Seal_2019-pool-5 | Juv | F | G |  | 5 |  |  |  |  |  |  |  |  |
| Seal_2019-pool-6 | Ad | F | G |  | 30 |  |  |  | 26 |  |  |  |  |
| Seal_2019-pool-7 | Ad | M | R |  | 10 |  |  |  | 17 |  |  |  |  |
| Seal_2019-pool-8 | Juv | F | R |  | 12 |  |  |  |  |  |  |  |  |
| Seal_2019-pool-9 | Ad | F | R |  | 393 |  |  |  | 33 |  |  |  |  |
| Seal_2020-pool-1 | Juv | M | C+B | 84 | 6 | 14 | 1 |  |  |  |  |  |  |
| Seal_2020-pool-2 | Juv | M | N | 32 |  | 10 |  |  |  |  |  |  | 5 |
| Seal_2020-pool-3 | Juv | M | G | 405 |  | 14 |  |  |  |  |  |  | 15 |
| Seal_2020-pool-4 | Juv | M | R | 121 |  | 852 |  |  |  |  |  | 1 | 29 |
| Seal_2020-pool-5 | Ad | M | C+B | 132 | 5 | 6 | 248 |  |  |  |  |  |  |
| Seal_2020-pool-6 | Ad | M | N | 16 |  | 1 | 17 |  |  |  |  |  |  |
| Seal_2020-pool-7 | Ad | M | G | 99 |  |  | 1 |  |  | 29 | 10 |  |  |
| Seal_2020-pool-8 | Ad | M | R | 18 |  |  | 15 |  |  |  |  |  |  |
| Seal_2020-pool-9 | Juv | F | C+B | 19 |  | 8 | 1 |  |  |  |  |  |  |
| Seal_2020-pool-10 | Juv | F | N | 268 |  | 22 | 4 |  |  |  |  |  |  |
| Seal_2020-pool-11 | Juv | F | G | 289 |  | 1 |  |  |  |  |  |  |  |
| Seal_2020-pool-12 | Juv | F | R | 50 |  |  |  |  |  | 11 |  | 1 |  |
| Seal_2020-pool-13 | Ad | F | C+B | 75 |  |  | 3 |  |  |  |  |  |  |
| Seal_2020-pool-14 | Ad | F | N |  | 8 |  | 34 |  |  |  |  |  |  |
| Seal_2020-pool-16 | Ad | F | R | 293 |  | 251 |  |  |  |  |  |  |  |
| Seal_2020-Dagestan* | Ad | F | SP |  |  |  | 1 |  |  |  |  | 1 |  |
| Total Contigs |  |  |  | 1945 | 556 | 1179 | 325 | 109 | 87 | 40 | 10 | 8 | 49 |

\*Sample from dead animal; Age: Ad – Adult, Juv – juvenile; Sex: F – female, M – male; Sample Type: B – buccal, N – nasal, G – genital, R – rectal, C – conjunctival, SP – spleen.

### Metagenomic sequencing and bioinformatics

For massively parallel sequencing, libraries were constructed using the NEBNext Ultra DNA Library Preparation kit (New England Biolabs, USA) according to the manufacturer’s protocol. Library size selection was performed using Ampure XP beads (Beckman Coulter, USA). The size and quality of libraries were checked on a Bioanalyzer 2100 instrument (Agilent Technologies, Germany). Sequencing was performed using the MiSeq Reagent version 3 kit on a MiSeq sequencer (Illumina, USA), at the Research and Production Center for Microbiology and Virology, Almaty, Kazakhstan. The reads were trimmed, and their quality was assessed with FastQC [40]. Sequence reads passing quality control filters were de novo assembled using Geneious 11.0 software (Biomatters, New Zealand) [41] by applying the installed SPAdes assembler [42] with default parameters. The resulting contigs were subjected to BLASTn and BLASTx searches in the local viral reference database as described in the Metavisitor pipeline [43]. Local BLAST hits with lengths >200 nucleotides (nt) were considered significant at E value <10^e-5^, and the potential viral sequences were subjected to a new BLASTx search against non-redundant protein sequences (nr) from the NCBI database. The best-hit sequences were individually annotated to the matching viral sequences. The viral sequences were verified by mapping reads to the corresponding reference genomes in the Geneious 11.0 software (Biomatters, New Zealand).

### PCR confirmation of viral detections

Following identification of viral contigs in the metagenomic analysis, PCR was carried out with primer sets synthesized for each viral species to confirm the presence of virus-specific nucleic acids in samples. PCR was undertaken using an Eppendorf Gradient amplifier using appropriate thermal cycling conditions. The presence of found viral sequences was verified using PCR/RT-PCR on the original RNA/DNA samples with corresponding primer sets [44-48].

*Phylogenetic analysis of metagenomic sequences and assessment of heterogeneity in virome diversity* Phylogenetic analysis of viral contigs derived from metagenomic sequencing, together with appropriate representative reference sequences, was carried out using the neighbor-joining method with 1000 bootstrap replicates using the p-distance in MEGA X [49].

Variation in sample viral diversity in relation to sample type, year, and sex-age class was visualized using a heatmap and cluster analysis implemented using the ComplexHeatmap package [50] in RStudio (v. 2022.12.0+353). The heatmap illustrates the virome diversity quantified by the number of contigs observed for each virus family within a particular pool/sample, where each original value has been incremented by 5 to enhance visualization clarity. A hierarchical cluster analysis was conducted in relation to sample type, by calculating a Euclidean distance based on viral abundances in each sample/pool, and then using the resulting distance matrix to cluster according the complete linkage algorithm.

## Results

### Metagenomic sequencing, BLAST search, and viral family relative abundance

An average of approximately 1,250,000 raw sequencing reads per library were obtained. Following assembly of contigs, BLAST searches revealed the presence of viruses belonging to 10 families, with viral contig counts ranging from 0 to 405 for each virus family-library combination (Table 1). Viral contigs ranged in size from 71 to 790 bp, and showed sequence identity ranging from 58% to 98.23% with known viruses (Table 2), suggesting some of these sequences might be derived from novel viral species.

**Table 2.** The most significant BLASTx hits matching known eukaryotic viruses, for contigs, obtained from Caspian seal samples.

| Family/Genus | Accession number | Contig length (nt) | Genome | Product | Best hit | Amino acid identity (%) | E-value | (RT)-PCR Confirmation |
| --- | --- | --- | --- | --- | --- | --- | --- | --- |
| *Circoviridae* | PP744471 | 207 | ssDNA | replication initiation protein | Replication initiation protein [Humpback whale blow-associated circo-like virus 2] (AWR89666) | 78.26 | 2e-29 | no |
| *Parvoviridae* | PP744472 | 175 | dsDNA | NS2 protein | NS2 protein [uncultured sea star-associated densovirus] (QOD39598) | 94.64 | 8e-28 | no |
| *Parvoviridae* | PP744473 | 144 | ssDNA | capsid protein VP1 | capsid protein VP1 [Canine parvovirus] (QBO24456) | 58.33 | 6e-09 | no |
| *Herpesviridae* | PP744474 | 312 | dsDNA | EUL25 protein | protein EUL25 [Equid alphaherpesvirus 1] (AII80971) | 81.33 | 1e-29 | yes |
| *Papillomaviridae* | PP744475 | 186 | dsDNA | E2 Protein | E2 protein [Leptonychotes weddellii papillomavirus 6] (NC_040818) | 72.58 | 2e-20 | no |
| *Picornaviridae* | PP744476 | 790 | +ssRNA | polyprotein | Polyprotein [Raccoon dog picornavirus] (UMO75562) | 72.59 | 2e-128 | no |
| *Caliciviridae* | PP744477 | 273 | +ssRNA | ORF-1 | ORF-1 protein [San Miguel sea lion virus 8] (AKG26793) | 63.74 | 5e-31 | no |
| *Anelloviridae* | PP744478 | 102 | ssDNA | ORF1 | ORF1 [Seal anellovirus 4] (YP_009115496.1) | 78.79 | 3e-09 | no |
| *Smacoviridae* | PP744479 | 96 | ssDNA | eplication associated protein | capsid protein [Swine associated smacovirus] (QCC72666) | 71.88 | 1e-04 | no |
| *Cruciviridae* | - | 183 | ssDNA | REP | putative replication associated protein [Crucivirus sp.] (QKV51008) | 45.59 | 4e-08 | no |
| *Orthomyxoviridae/ Influenza A* | PP744480 | 71 | -ssRNA | hemagglutinin gene | hemagglutinin gene (Influenza A virus A/Anas platyrhynchos/Belgium/3950-8/2015(H3N8)) [MT406985] | 95.83 | 1e-21 | yes |
| *Orthomyxoviridae/ Influenza A* | PP744481 | 276 | -ssRNA | nonstructural protein | nonstructural protein 1 (NS1) gene [Influenza A virus A/Blue-winged Teal/Kansas/AH0029699S.8.B/2015 (H12N6)] (QDX48107) | 92.94 | 3e-48 | yes |
| *Orthomyxoviridae/ Influenza B* | PP744482 | 312 | -ssRNA | polymerase PB2 | polymerase PB2 [Influenza B virus (B/Florida/09/2016)] (APW80077) | 97.12 | 4e-59 | yes |
| *Orthomyxoviridae/ Influenza B* | PP744483 | 360 | -ssRNA | NS protein | NS protein [Influenza B virus (B/Arizona/38/2016)] (AQS98133) | 98.23 | 7e-75 | yes |

Overall, the most abundant viral contigs (Figure 1.), comprising 45% of those recovered, matched the family *Circoviridae*. More than a quarter (27.3%) were assigned to parvoviruses, (12.9%) matched herpesviruses, followed by *Papillomaviridae* (7.5%), *Picornaviridae* (2.5%), *Caliciviridae* (2%), *Anelloviridae* and *Cruciviridae* (both ∼1%). The rarest contigs belonged to *Smacoviridae* (0.23%) and *Orthomyxoviridae* (0.16%).

**Figure 1.**
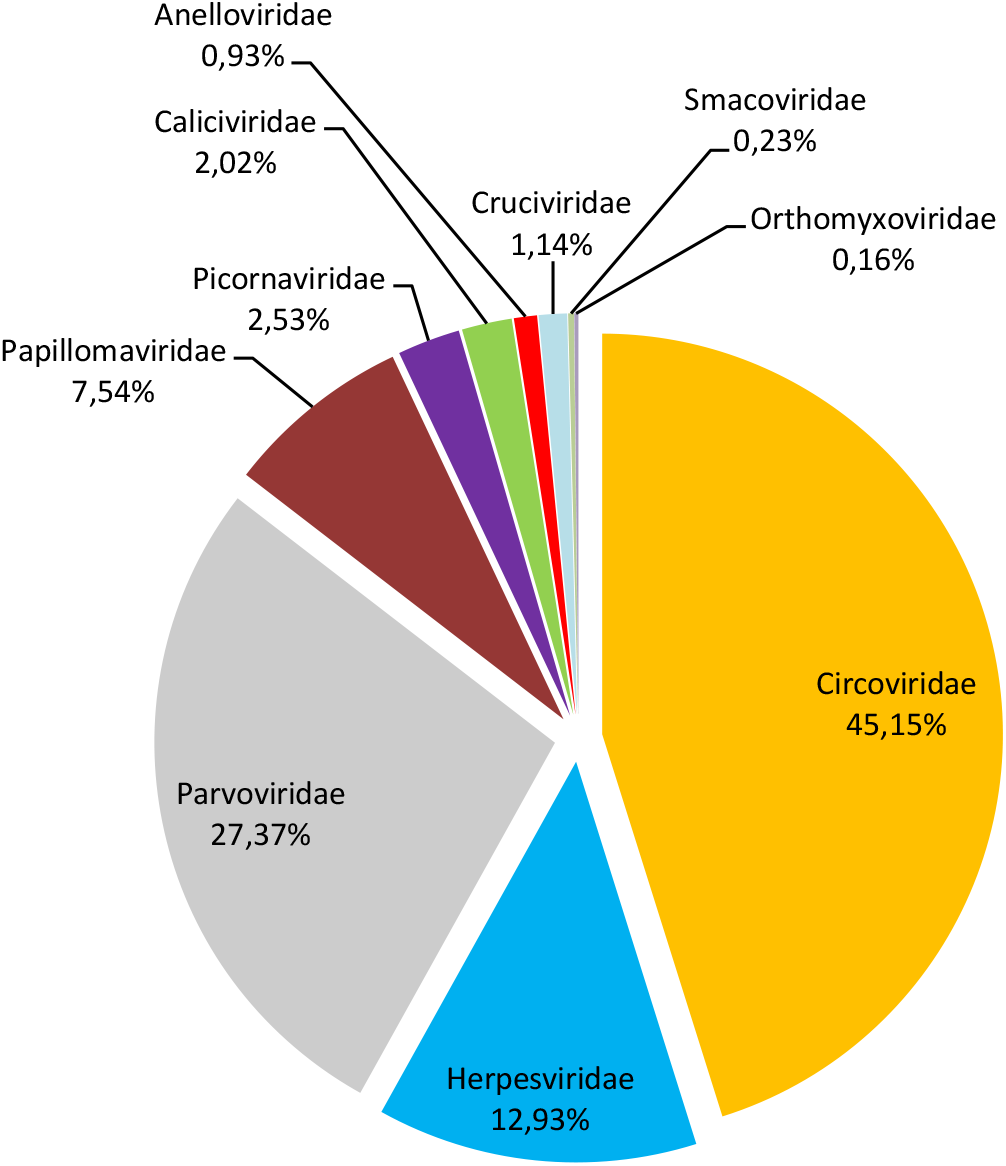
Proportions of viral families in across all samples

### PCR confirmation

Of the 10 viral families found in metagenomic sequencing, confirmatory PCR amplicons were recovered only for viruses from *Herpesviridae* and *Orthomyxoviridae*. Additionally, although no *Adenovirus* contigs were recovered in the metagenomic sequencing, PCR-screening detected adenovirus products from buccal, nasal and rectal swabs from six live seals sampled in 2020, adding to PCR detections of adenovirus in samples from earlier years reported in our previous study [33].

Previously we reported molecular and serological evidence for ongoing circulation of Canine morbillivirus (formerly canine distemper virus; CDV), a member the family *Paramyxoviridae*, in Caspian seals [33]. No CDV contigs were detected for metagenomic sequencing samples collected 2009-2020, and RT-PCR testing for CDV in 2019-2020 samples was also negative.

### Heterogeneity in sample virome diversity

Analysis of the age-specific virus patterns revealed a notable disparity in the prevalence of circovirus and parvovirus contigs, with juvenile seals exhibiting a higher abundance compared to adults, a trend that was particularly pronounced in 2020. Additionally, the results highlight the high prevalence of herpesvirus among adult seals in 2019. *Parvoviridae, Papillomaviridae, Anelloviridae, Smacoviridae*, and *Cruciviridae* families were found only in samples from 2020, with *Caliciviridae* family found only in 2019, and *Picornaviridae* present only in 2016. The majority of the *Herpesviridae* sequences were recovered from 2019 samples, but were also found at low frequency in 2016 and 2020. *Circoviridae* contigs were overwhelmingly found in 2020, had smaller numbers 2016, and were absent in 2019 (Table 1., Figure 2).

**Figure 2.**
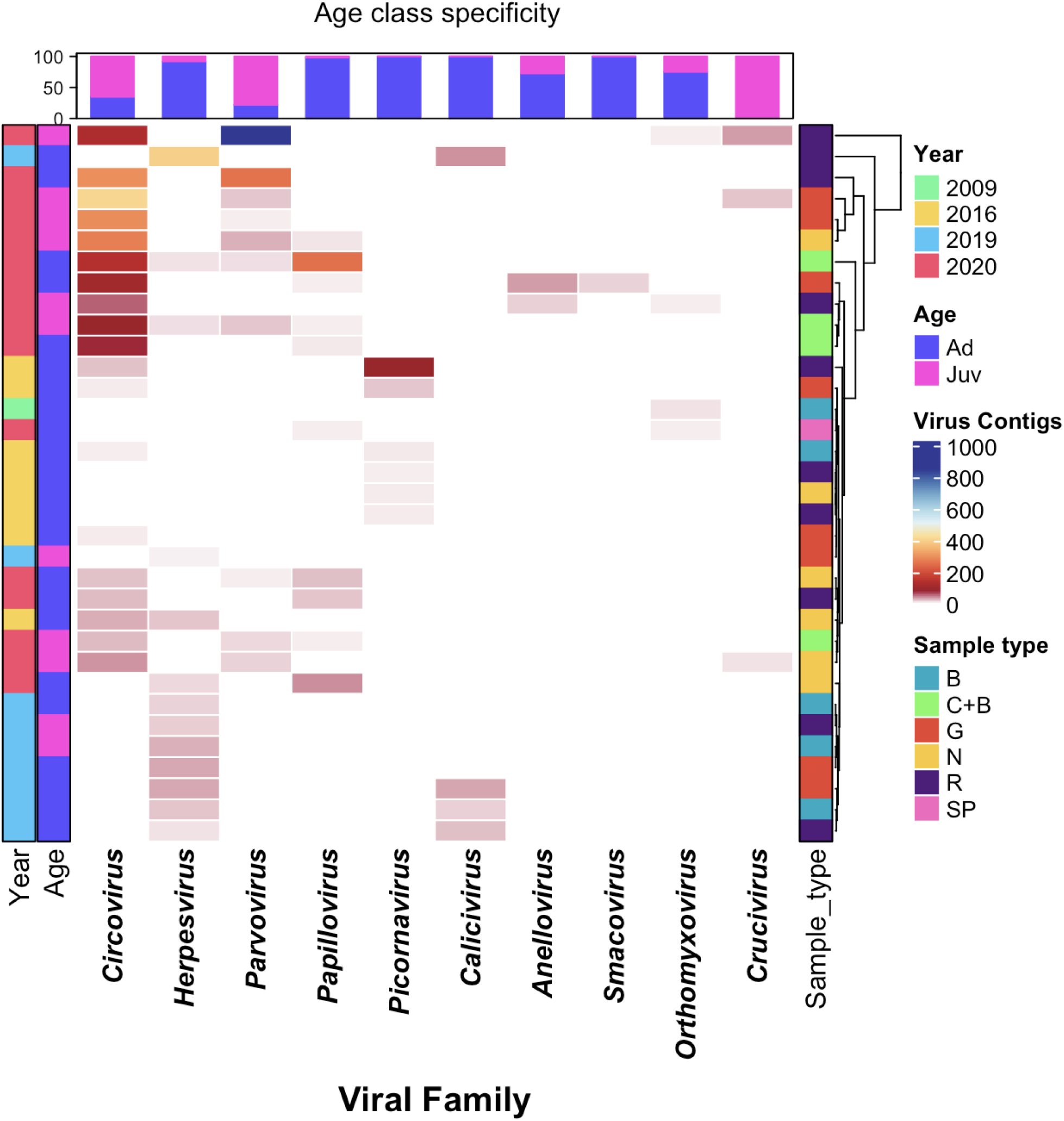
Heatmap of virome diversity of seals, based on Age and Sample types across different years, illustrating virus relative abundance profiles, with each cell representing the contig counts belonging to the given virus family, indicated by corresponding colors. Samples are from dead seals; Age: Ad – Adult, Juv – Juvenile; Sample Type: B – Buccal, N – Nasal, G – Genital, R – Rectal, C – Conjunctival, SP – Spleen

### Description of identified viral contigs

The *Circoviridae* is a family of small non-enveloped circular single-stranded DNA viruses identified from various mammals, birds, and other organisms worldwide. A single contig type of 207 nt was recovered, matching the replication initiation protein from humpback whale blow-associated Circo-like virus 2, discovered in Australia in 2017 [51], with a BLASTx amino acid similarity of 78.26%. Phylogenetic analysis (Supplementary Figure 1A) supported the same association. The second most abundant family is *Parvoviridae*, which are non-enveloped linear ssDNA viruses. Two distinct parvovirus contig were found. The first, matching Parvovirus-like VP1 gene sequences, was 58.33% similar to Canine parvovirus in BLASTx searches. The second, matching Parvovirus-like nonstructural NS2 gene sequences, was 94.64% similar to densoviruses found in starfish, invertebrates, insects, and decapod crustaceans [52]. Phylogenetic analyses with the partial NS2 protein sequence (Supplementary Figure 1B) suggest they are associated with parvoviruses from other aquatic organisms, clustering with densoviruses from oyster, mollusks, and sea stars.

*Herpesviridae* is a large family of enveloped dsDNA viruses [1]. Our previous study found partial Us2-like protein sequences in Caspian seals that were 100% similar to Phocid alphaherpesvirus 1 [33] described in Atlantic harbor seals in 1985 [53]. In this study contigs were recovered for a fragment of the EUL25-like protein gene with 81.33% similarity to Equid alphaherpesvirus 1 in BLASTx analysis, which formed a separate cluster from the corresponding Phocid herpesvirus 1 cluster in a phylogenetic tree (Supplementary Figure 1C).

*Papillomaviruses* are small, non-enveloped, circular dsDNA viruses that infect skin or mucosal membranes in a wide range of animals, including marine mammals [54]. Caspian seals had sequences for one contig that was 72.58 % similar to the E2 protein sequence of papillomavirus 6 from Weddell seals in BLASTx searches [22]. Phylogenetically, the Caspian seal papillomavirus clusters most closely with other mammalian Papillomaviruses from rodents, mustelids and cats. (Supplementary Figure 1D).

*Picornaviruses* are small, non-enveloped, positive sense ssRNA viruses. From Caspian seals, we recovered a contig of 790 nt with 72.59 % similarity in BLASTx analysis to a Racoon dog picornavirus polyprotein gene protein isolated in China in 2017-2021. Phylogenetic analysis suggested that Caspian seal picornavirus was taxonomically closest to California sea lion sapelovirus, with both forming a clade as part of a lineage also containing Racoon dog picornavirus, and Feline picornavirus (Supplementary Figure 1E).

*Caliciviridae* is a non-enveloped single-stranded, positive sense RNA viral family. Marine caliciviruses belong to the genus *Vesivirus* [55]. It has been suggested that marine mammals serve as their natural reservoir [56]. In BLASTx searches, a 273 nt contig for vesivirus-like sequences found in this study were 63.74% similar to the ORF-1 protein sequence from San Miguel sea lion virus 8, isolated in the USA [57].

Phylogenetic analyses confirmed its relatedness to the San Miguel sea lion virus and canine vesiviruses (Supplementary Figure 1F).

*Anelloviruses* are non-enveloped, circular ssDNA viruses belonging to the *Anelloviridae* family [18]. They are highly diverse and to date have not been associated with any clinically manifesting disease [58]. The Caspian seal anellovirus sequence, represented by a 102 nt contig, was 78.79% similar in BLASTx search to Seal anellovirus 4 ORF-1 protein [25]. Phylogenetic analysis of the partial ORF1 nucleotide sequence of Caspian seal anellovirus showed its relatedness to a cluster consisting of Seal anelloviruses 4 and 5 (Supplementary Figure 1G).

*Smacoviridae and Cruciviridae* family sequences were found in samples from 2020. They both possess small circular single-stranded DNA genomes. S*macoviridae* is frequently found in vertebrate species but not in environmental samples and was previously known as “stool-associated circular viruses” detected in healthy and diarrheic mammals, birds as well as in insect species [59], with one previous report in sea lions [10]. In BLASTx searches, the Caspian seal 96 nt smacovirus contig sequence was 71.88% similar to Swine associated smacovirus capsid protein. *Cruciviridae* possess chimeric genome that encompass genes derived from eukaryotic ssRNA and ssDNA viruses [60]. The putative crucivirus 189 nt sequence in BLASTx search was 45.59% similar to replication associated protein from crucivirus found during metagenomic data mining of environmental water [61].

*Orthomyxoviruses* are enveloped viruses with a segmented RNA genome. We previously reported Influenza A virus RNA detected by RT-PCR in samples from Caspian seals collected in 2009 [33] but their sequences were not determined at the time. Here, two contig types were recovered from one of the 2009 samples, for hemagglutinin and nonstructural protein genes of Influenza A virus (Table 2). BLASTx searches revealed they were 95.8% and 92.94% similar to influenza virus sequences of H3 and H12 subtypes isolated from wild birds in 2006 - 2019 worldwide. Phylogenetic analysis of the H3 hemagglutinin and non-structural protein genes confirmed the relatedness of the Caspian influenza A sequences to the avian Eurasian lineage (Supplementary Figures 1H and 1I).

Previously influenza B virus, a predominantly human associated virus, has described in harbour and grey seals (*Halichoreus grypus*) [47], and serological evidence has been found for its circulation in Caspian seals [33, 62]. Here, two distinct influenza B virus contigs were identified, for the first time in Caspian seals, in the spleen sample from the dead animal stranded in Dagestan (Russia) in 2020. One sequence matched the PB2 polymerase gene, showing 97.12% similarity with the B/Florida/09/2016 virus isolated from humans. The second from the nonstructural NS gene, showed 98.23% similarity with the B/Arizona/38/2016 virus. Phylogenetic analyses returned a similar pattern of clustering (Supplementary Figures 1J and 1K).

## Discussion

In this study, we made the first evaluation of viral diversity of Caspian seals using a metagenomic approach, recovering viral contigs belonging to families whose representatives are linked to both apathogenic or disease-causing infections in mammals. The most abundant contigs observed in samples belonged to the *Circoviridae* family. Pathogenic circovirus infections are found in porcine and avian species worldwide. Subclinical infections are more common, but porcine circovirus 1, infectious chicken anemia, and psittacine beak and feather disease are associated with the severe disease course [63]. The Caspian seal circovirus was 78.26% similar to the recently identified novel circovirus-like virus detected in Humpback whale blow [51]. These single stranded DNA circo-like viruses appear to be significant virome components in many marine environments and marine invertebrates may be the primary host [64]. For now, the biological and clinical significance of these novel circovirus for marine mammals remains unknown.

*Parvoviridae* was the second most abundant family in tested samples. Densoviruses are known to infect invertebrates, insects, and decapod crustaceans [52]. Thus, the Parvovirus-like sequences found in Caspian seals appear related to parvoviruses associated with aquatic ecosystems. These findings parallel those for a study of the humpback whale blow virome where sequences from *Circoviridae* and *Parvoviridae* families also formed the top two most abundant read classes [51].

In the *Herpesviridae* family, to date seven phocid herpesviruses have been recorded worldwide, phocid herpesviruses from type-1 to type-7 (PhHV 1 - 7) [25]. Newborn and immunosuppressed seals infected with PhHV-1 die with symptoms of acute pneumonia and focal hepatitis but adults show no apparent infection [46]. PhHV-2 is more distantly related to both PhHV-1 and herpesviruses of terrestrial carnivores [46]. Previously, we found a herpesvirus Us2-like protein partial sequence in Caspian seals [33] that was 100% similar to Phocid alphaherpesvirus 1 from Atlantic harbor seal [53]. In the current study we recovered partial contigs for the EUL25 protein with 81.33% similarity to Equid alphaherpesvirus 1. Further studies are necessary to obtain the complete genome of the Phocid alphaherpesvirus 1 from Caspian seals for full characterization, and to determine the number of strains. If PhHV-1 from Caspian seal behaves similarly to other alphaherpesviruses, it may establish latency in neurons [65], without manifesting any clinical signs, but additional work is needed to determine infection mechanisms.

In marine mammals, *Papillomaviridae* family viruses have been found in manatees [66], cetaceans [67] and pinnipeds [68]. Clinical signs include warty lesions in lingual and genital mucosa and cutaneous tissues. Some papillomavirus species have been identified as oncogenic [65]. Papillomavirus from Caspian seals by E2 protein was 72.58% similar to *Leptonychotes weddellii* papillomavirus 6, but proliferative axillary lesions were not observed in Caspian seals examined or probably were not visible.

*Picornaviridae* family sequences found in samples from Caspian seals were 72.59% similar to Racoon dog picornavirus isolated from healthy animals in China in 2021 [69]. However, phylogenetic analysis indicated that Caspian seal picornavirus was clustered closest to California sea lion sapelovirus in a lineage that also include the Racoon dog virus. Raccoon dogs are widely spread in the Northern Caspian seashore and frequently visit haul-out sites of Caspian seals. So, it is not excluded that this picornavirus species circulates in local carnivore and pinniped populations. Picornaviruses cause notable veterinary diseases with significant mortality, including foot-and-mouth disease in hoofed animals and avian encephalomyelitis in poultry. In marine mammals, picornaviruses have been found in sub-Antarctic fur seals, South American fur seals [18], California sea lions [16], harbor seals [70], ribbon and ringed seals [71], but their role in pathology still needs to be determined.

The *Caliciviridae* family was detected in samples from Caspian seals from fragments of nonstructural polyprotein gene sequences that were 63.74% similar to San Miguel sea lion virus (SMSV), identified in the USA in 1972. This San Miguel sea lion virus is a marine calicivirus that belongs to a distinct lineage within the genus *Vesivirus*, which also includes Vesicular Exanthema of Swine Virus (VESV). VESV is clinically indistinguishable from picornaviral vesicular diseases such as foot-and-mouth disease, swine vesicular disease and rhabdoviral vesicular stomatitis. It has been suggested that VESV, SMSV, and other related viruses originated from marine mammal caliciviruses [57], with marine vesiviruses circulating worldwide. Clinically, SMSV causes vesicular lesions on the flippers and the mouth, as well as gastroenteritis in California sea lions [57]. No lesions were observed in Caspian seals at the time of sampling, although the missing of small lesions cannot be ruled out.

*Anelloviridae* sequences detected in Caspian seals were 78.79% similar to ORF1 of Seal anellovirus 4 identified from harbor seals (*Phoca vitulina*) in the Netherlands in 2014. The role of anelloviruses in the pathology of wild animals, including pinnipeds, is not well understood, although many viruses have been genetically characterized. For human anelloviruses, a clear link between anellovirus positivity and disease has not been established [72], but recent studies have suggested that some anelloviruses are associated with unexplained fever, diabetes, cirrhosis in liver transplant patients, respiratory disease, cancer, and autoimmune disorders [58].

Influenza viruses of the *Orthomyxoviridae* family are also significant for the health status of marine mammals. Diseases caused by influenza viruses of avian origin from subtypes H7N7 [73], H4N5 [74], H3N3 [75] and H13 [76] subtypes are known in pinnipeds. A pandemic pdmH1N1 strain [77], that spread globally in 2009 has also been isolated from healthy-looking seals, which indicates the potential of seals to serve as a reservoir of influenza viruses in the wild. Additionally, a H5N1 has recently spilled over into pinnipeds in both the northern and southern hemispheres [13, 14, 78], [79] and in Antarctica [80] causing mass mortalities.

In this study, we make the first report of sequence of the H3 influenza A hemagglutinin gene to be detected from Caspian seals as well as the nonstructural protein. Previously, H4 influenza A subtype was described in Caspian seals [81]. This hemagglutinin sequence was 95.83% similar to H3 subtype influenza viruses of the Eurasian lineage isolated from wild birds. This subtype was also previously found in pinnipeds [75]. The sequence was derived from a sample collected in 2009 from a live, apparently healthy individual. Antibodies to the influenza A virus have previously been found in the Caspian seal [33, 62] with RT-PCR confirmation [33], but to date influenza A has not been linked conclusively to significant mortality events in Caspian seals. Analysis of marine mammal influenza viruses suggests they were acquired through close contact with wild birds such as oral–fecal transmission through seawater [74]. In the Caspian region, sea and migratory birds regularly share seal haul out sites, creating opportunities for this route of transmission [33, 82]. Consumption of carcasses of dead birds is not part of Caspian seals’ diet and was never reported before [26, 78, 83], so infection with influenza viruses through this route, as has been suggested for some recent H5N1 transmissions elsewhere [84], may be less likely.

Influenza B virus sequences were also identified in Caspian seals. Influenza B viruses were first isolated from a juvenile harbour seal in a rehabilitation center in the Netherlands [47]. It was suggested that the virus had been introduced in the seal population from a human source and circulated in seals for some time. Further, antibodies to influenza B were detected in harbour and grey seals in the 2010s [85], also in Caspian seals [33, 61]. In this study, influenza B sequences were found in a spleen sample from a dead seal that stranded on the in the Dagestan seashore during a mass mortality event in 2020 [38]. BLAST search has shown that the partial sequences of the Caspian seal influenza B virus PB2 and NS genes were closely related to those of strains circulating in humans simultaneously in 2020 and five years earlier (data not shown). However, this finding in isolation is not sufficient to attribute influenza B as factor in the mass mortality. At this time there is not sufficient data to consider seals a reservoir of influenza B in nature, but we can see that seals are also susceptible to this virus and the routes of seal exposure are still to be determined.

*Smacoviridae and Cruciviridae* are viral families that have not been studied in marine mammals and their evolutionary and ecological history in this context is unclear. At this time, we can only state their presence in the viral metagenome of the Caspian seal, and further research is needed to deepen our knowledge of these viral families.

*Adenoviridae* family viruses are double-stranded DNA viruses that infect all groups of vertebrates, including marine mammals. Canine adenovirus 1 causes viral hepatitis in dogs and infects many wildlife species [86]. Adenoviruses are known to cause fatal hepatitis in different otariid species [45, 87]. In our previous research, we have identified an adenovirus sequence that is 76 % [33] similar to Phocine adenovirus 1 from a Northern elephant seal (*Mirounga angustirostris*) with ocular lesions [88]. It was suggested that the presence of this virus in ocular tissues of pinnipeds was common and not significantly associated with the disease [88]. Here we have not detected adenoviral sequences from metagenomic sequencing, but PCR screening revealed positive samples collected from live seals in 2020, adding to those from our earlier study. This suggests that direct PCR detection may be more sensitive than NGS based approaches when testing swab samples from seals.

Representatives of the genus *Morbillivirus* of the *Paramyxoviridae* family are some of the most important pathogens for marine mammals. CDV has previously caused mass mortalities of thousands of animals in Caspian and Baikal seals (*Pusa sibirica*) [31, 89]. Our previous work detected antibodies to CDV in Caspian seals with ELISA tests for six sera out of 74 (8.1%) animals sampled from 2007 to 2017, and 5 of 13 asymptomatic animals sampled 2008 were positive for PCR tests [33]. Two complete CDV genomes were recovered for the 2008 animals, which formed a clade with 99.59% identity to the 2000 Caspian seal epizootic lineage [33]. In the current work, no morbillivirus contigs were recovered from metagenomic sequencing of samples 2009-2020, and the were no positive detections by RT-PCR for 2019-2020 samples. Together, our two studies and others [90], suggest that CDV has circulated episodically in Caspian seals since 2001, but the trigger mechanisms for mass mortalities are still unknown.

Our study has some limitations. Firstly, only short viral sequences were identified in the metagenomic sequencing with a maximum contig size of 790 nt. These short sequence lengths mean that the phylogenetic position of the viruses requires further confirmation, but this is also restricted by reference sequence coverage for most of the families. Full viral genomes will greatly enhance the capacity to determine viral identity, phylogenetic relationships and evaluations of disease potential, and functional diversity. Secondly, it was not possible to independently confirm viral presence via PCR/RT-PCR in all cases due to lack of reliable conserved primers for all families. Expanding genomic coverage for these viruses will support future primer development for use in diagnostic and confirmatory testing. Lastly, inferring viral abundance from contig read counts is an indirect indicator, and should be interpreted cautiously, but is commonly taken as representative of relative prevalence in metagenomic studies [15, 16, 18].

In conclusion, the viral metagenomic profile of Caspian seals is similar to those from other marine mammals and all viral families described in this research have previously been reported in marine mammals. The two most dominant viral families consisting of *Circoviridae* and *Parvoviridae* families found in this study are widespread in marine ecosystems. The primary hosts of these viruses are likely to be Caspian Sea marine invertebrates in the case of circoviruses, or insects and crustaceans in the case of densoviruses in the *Parvoviridae* family. These two families make up 72% of the entire Caspian seal virome, and are possibly of dietary origin, since Caspian seals frequently feed on crustaceans and invertebrate consuming fish [83].

A second major group in the virome comprises mammalian viruses: *Herpesviridae, Papillomaviridae, Caliciviridae, Anelloviridae, Adenoviridae, Orthomyxoviridae* and *Paramyxoviridae*. Viruses of this group can potentially cause various pathologies in mammals or be asymptomatic [91]. It is still unclear to which group the *Picornaviridae*-like sequences belong since the identified contigs are genetically distant from mammalian and aquatic viruses. The similarity of the Caspian seal viruses of both groups with those registered in other animals worldwide varied from 58 to 100%. The habitat of the Caspian seal is restricted to the landlocked Caspian Sea, with no direct access to the World Ocean. The Caspian Sea is a remnant of the Paratethys, and is last thought to have shared an open connection the world’s ocean 35 million years ago, although sporadic connections with the Arctic Ocean and Mediterranean Sea likely existed through Pleistocene glacial cycles [92, 93]. Caspian seals are estimated to have diverged from sister taxa around 1 to 2 million years ago [94]. Therefore, given this history of environmental and species isolation, the discovery of genetically distant viruses was expected. As virome data from marine mammals accumulates, results such as these can contribute to comparative studies of the ecology and evolution of pathogen communities and their implications for marine mammal health. These results will contribute to new resources and strategies for disease surveillance in Caspian seals. Given the regular occurrence of mass mortalities, with annual strandings hundreds to thousands of carcasses, it is essential to develop capacity for pathology investigations to support integrated Caspian seal conservation [33], with particular priority given to coordinated transboundary surveillance for influenza and CDV.

## Acknowledgments

The authors are grateful to Mart Jüssi, Timur Baimukanov, Lilia Dmitrieva, Mikhail Verevkin, Assel Baimukanova, Fedor Klimov, Alexey Mulyaev, Mirgaliy Baimukanov, and Ivar Jüssi for their help in sampling from seals.

