## Supplementary Figure S1 for "VIRAL METAGENOMIC SURVEY OF CASPIAN SEALS"

A.

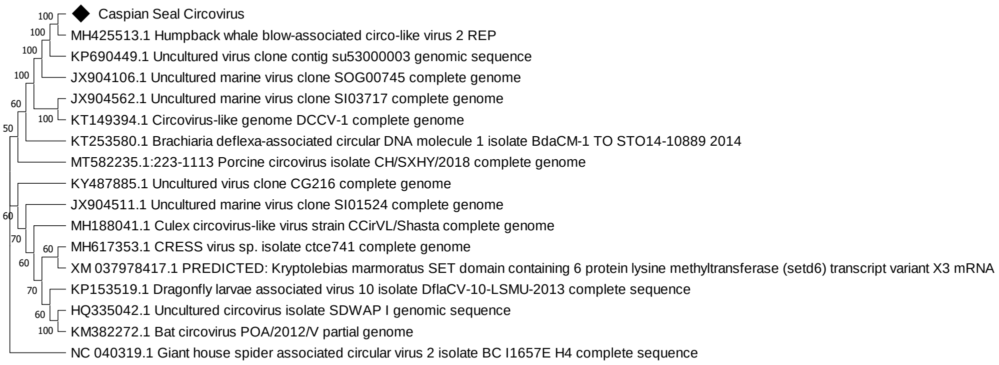
Caspian Seal Circovirus
MH425513.1 Humpback whale blow-associated circo-like virus 2 REP
KP690449.1 Uncultured virus clone contig su53000003 genomic sequence
JX904106.1 Uncultured marine virus clone SOG00745 complete genome
JX904562.1 Uncultured marine virus clone SI03717 complete genome
KT149394.1 Circovirus-like genome DCCV-1 complete genome
KT253580.1 Brachiaria deflexa-associated circular DNA molecule 1 isolate BdaCM-1 TO STO14-10889 2014
MT582235.1:223-1113 Porcine circovirus isolate CH/SXHY/2018 complete genome
KY487885.1 Uncultured virus clone CG216 complete genome
JX904511.1 Uncultured marine virus clone SI01524 complete genome
MH188041.1 Culex circovirus-like virus strain CCirVL/Shasta complete genome
MH617353.1 CRESS virus sp. isolate ctce741 complete genome
XM 037978417.1 PREDICTED: Kryptolebias marmoratus SET domain containing 6 protein lysine methyltransferase (setd6) transcript variant X3 mRNA
KP153519.1 Dragonfly larvae associated virus 10 isolate DflaCV-10-LSMU-2013 complete sequence
HQ335042.1 Uncultured circovirus isolate SDWAP I genomic sequence
KM382272.1 Bat circovirus POA/2012/V partial genome
NC 040319.1 Giant house spider associated circular virus 2 isolate BC I1657E H4 complete sequence

B.

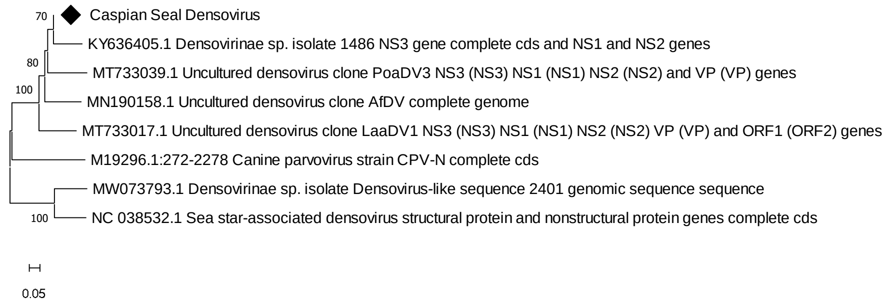
Caspian Seal Densovirus
KY636405.1 Densovirinae sp. isolate 1486 NS3 gene complete cds and NS1 and NS2 genes
MT733039.1 Uncultured densovirus clone PoaDV3 NS3 (NS3) NS1 (NS1) NS2 (NS2) and VP (VP) genes
MN190158.1 Uncultured densovirus clone AfDV complete genome
MT733017.1 Uncultured densovirus clone LaaDV1 NS3 (NS3) NS1 (NS1) NS2 (NS2) VP (VP) and ORF1 (ORF2) genes
M19296.1:272-2278 Canine parvovirus strain CPV-N complete cds
MW073793.1 Densovirinae sp. isolate Densovirus-like sequence 2401 genomic sequence sequence
NC 038532.1 Sea star-associated densovirus structural protein and nonstructural protein genes complete cds
0.05

C.

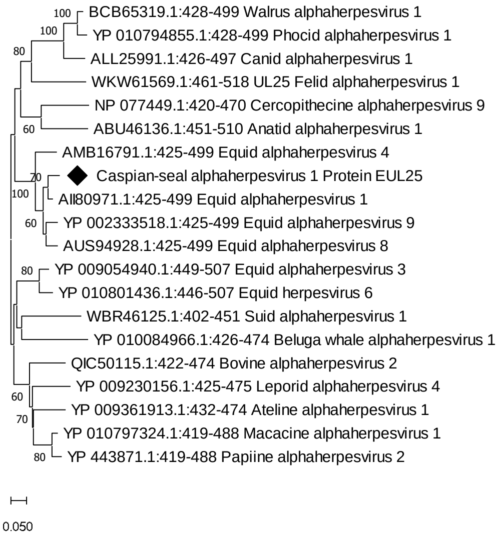
BCB65319.1:428-499 Walrus alphaherpesvirus 1
YP 010794855.1:428-499 Phocid alphaherpesvirus 1
ALL25991.1:426-497 Canid alphaherpesvirus 1
WKW61569.1:461-518 UL25 Felid alphaherpesvirus 1
NP 077449.1:420-470 Cercopithecine alphaherpesvirus 9
ABU46136.1:451-510 Anatid alphaherpesvirus 1
AMB16791.1:425-499 Equid alphaherpesvirus 4
Caspian-seal alphaherpesvirus 1 Protein EUL25
AII80971.1:425-499 Equid alphaherpesvirus 1
YP 002333518.1:425-499 Equid alphaherpesvirus 9
AUS94928.1:425-499 Equid alphaherpesvirus 8
YP 009054940.1:449-507 Equid alphaherpesvirus 3
YP 010801436.1:446-507 Equid herpesvirus 6
WBR46125.1:402-451 Suid alphaherpesvirus 1
YP 010084966.1:426-474 Beluga whale alphaherpesvirus 1
QIC50115.1:422-474 Bovine alphaherpesvirus 2
YP 009230156.1:425-475 Leporid alphaherpesvirus 4
YP 009361913.1:432-474 Ateline alphaherpesvirus 1
YP 010797324.1:419-488 Macacine alphaherpesvirus 1
YP 443871.1:419-488 Papiine alphaherpesvirus 2
0.050

D.

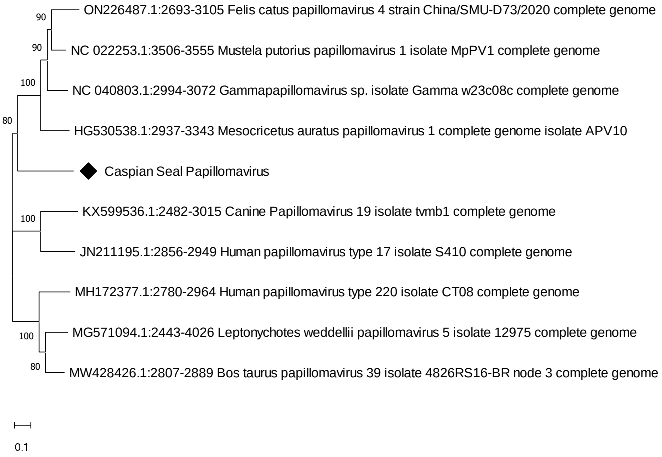
ON226487.1:2693-3105 Felis catus papillomavirus 4 strain China/SMU-D73/2020 complete genome
NC 022253.1:3506-3555 Mustela putorius papillomavirus 1 isolate MpPV1 complete genome
NC 040803.1:2994-3072 Gammapapillomavirus sp. isolate Gamma w23c08c complete genome
HG530538.1:2937-3343 Mesocricetus auratus papillomavirus 1 complete genome isolate APV10
Caspian Seal Papillomavirus
KX599536.1:2482-3015 Canine Papillomavirus 19 isolate tvmb1 complete genome
JN211195.1:2856-2949 Human papillomavirus type 17 isolate S410 complete genome
MH172377.1:2780-2964 Human papillomavirus type 220 isolate CT08 complete genome
MG571094.1:2443-4026 Leptonychotes weddellii papillomavirus 5 isolate 12975 complete genome
MW428426.1:2807-2889 Bos taurus papillomavirus 39 isolate 4826RS16-BR node 3 complete genome
0.1

E.

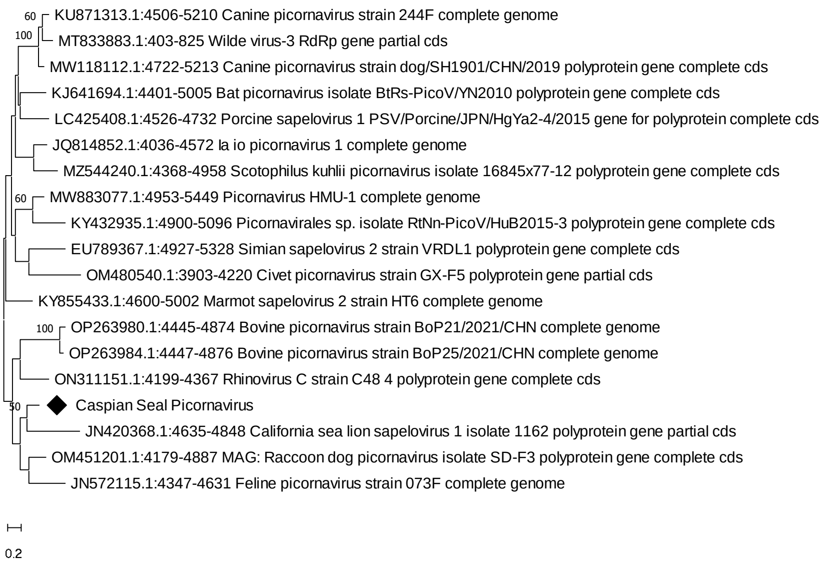
60
100
KU871313.1:4506-5210 Canine picornavirus strain 244F complete genome
MT833883.1:403-825 Wilde virus-3 RdRp gene partial cds
MW118112.1:4722-5213 Canine picornavirus strain dog/SH1901/CHN/2019 polyprotein gene complete cds
KJ641694.1:4401-5005 Bat picornavirus isolate BtRs-PicoV/YN2010 polyprotein gene complete cds
LC425408.1:4526-4732 Porcine sapelovirus 1 PSV/Porcine/JPN/HgYa2-4/2015 gene for polyprotein complete cds
JQ814852.1:4036-4572 la io picornavirus 1 complete genome
MZ544240.1:4368-4958 Scotophilus kuhlii picornavirus isolate 16845x77-12 polyprotein gene complete cds
60
MW883077.1:4953-5449 Picornavirus HMU-1 complete genome
KY432935.1:4900-5096 Picornavirales sp. isolate RtNn-PicoV/HuB2015-3 polyprotein gene complete cds
EU789367.1:4927-5328 Simian sapelovirus 2 strain VRDL1 polyprotein gene complete cds
OM480540.1:3903-4220 Civet picornavirus strain GX-F5 polyprotein gene partial cds
KY855433.1:4600-5002 Marmot sapelovirus 2 strain HT6 complete genome
100
OP263980.1:4445-4874 Bovine picornavirus strain BoP21/2021/CHN complete genome
OP263984.1:4447-4876 Bovine picornavirus strain BoP25/2021/CHN complete genome
ON311151.1:4199-4367 Rhinovirus C strain C48 4 polyprotein gene complete cds
Caspian Seal Picornavirus
50
JN420368.1:4635-4848 California sea lion sapelovirus 1 isolate 1162 polyprotein gene partial cds
OM451201.1:4179-4887 MAG: Raccoon dog picornavirus isolate SD-F3 polyprotein gene complete cds
JN572115.1:4347-4631 Feline picornavirus strain 073F complete genome
0.2

F.

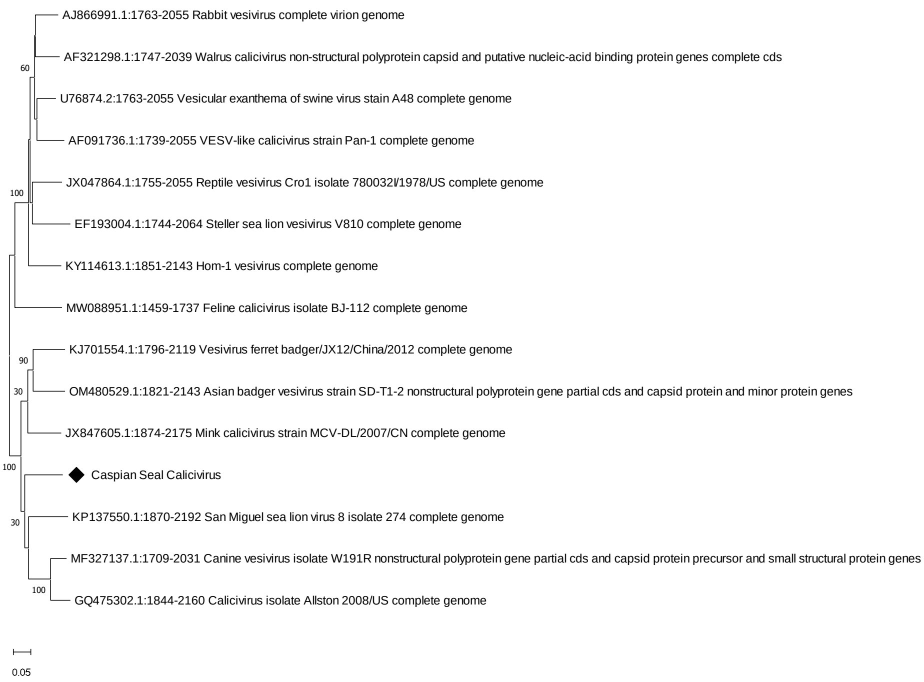
AJ866991.1:1763-2055 Rabbit vesivirus complete virion genome
AF321298.1:1747-2039 Walrus calicivirus non-structural polyprotein capsid and putative nucleic-acid binding protein genes complete cds
60
U76874.2:1763-2055 Vesicular exanthema of swine virus stain A48 complete genome
AF091736.1:1739-2055 VESV-like calicivirus strain Pan-1 complete genome
JX047864.1:1755-2055 Reptile vesivirus Cro1 isolate 780032I/1978/US complete genome
100
EF193004.1:1744-2064 Steller sea lion vesivirus V810 complete genome
KY114613.1:1851-2143 Hom-1 vesivirus complete genome
MW088951.1:1459-1737 Feline calicivirus isolate BJ-112 complete genome
KJ701554.1:1796-2119 Vesivirus ferret badger/JX12/China/2012 complete genome
90
OM480529.1:1821-2143 Asian badger vesivirus strain SD-T1-2 nonstructural polyprotein gene partial cds and capsid protein and minor protein genes
30
JX847605.1:1874-2175 Mink calicivirus strain MCV-DL/2007/CN complete genome
100
Caspian Seal Calicivirus
KP137550.1:1870-2192 San Miguel sea lion virus 8 isolate 274 complete genome
30
MF327137.1:1709-2031 Canine vesivirus isolate W191R nonstructural polyprotein gene partial cds and capsid protein precursor and small structural protein genes
100
GQ475302.1:1844-2160 Calicivirus isolate Allston 2008/US complete genome
0.05

G.

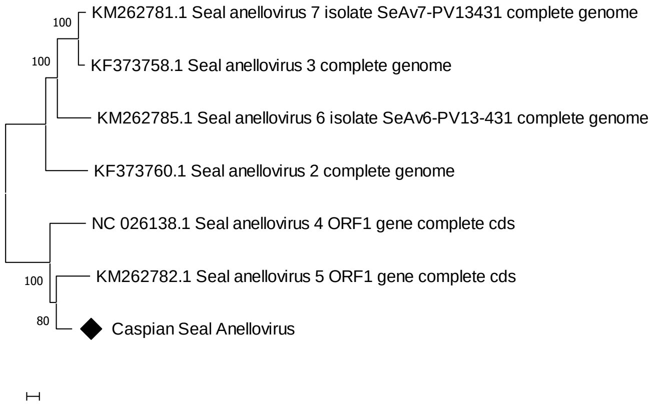
100
KM262781.1 Seal anellovirus 7 isolate SeAv7-PV13431 complete genome
100
KF373758.1 Seal anellovirus 3 complete genome
KM262785.1 Seal anellovirus 6 isolate SeAv6-PV13-431 complete genome
KF373760.1 Seal anellovirus 2 complete genome
NC 026138.1 Seal anellovirus 4 ORF1 gene complete cds
100
KM262782.1 Seal anellovirus 5 ORF1 gene complete cds
80
Caspian Seal Anellovirus
0.05

H.

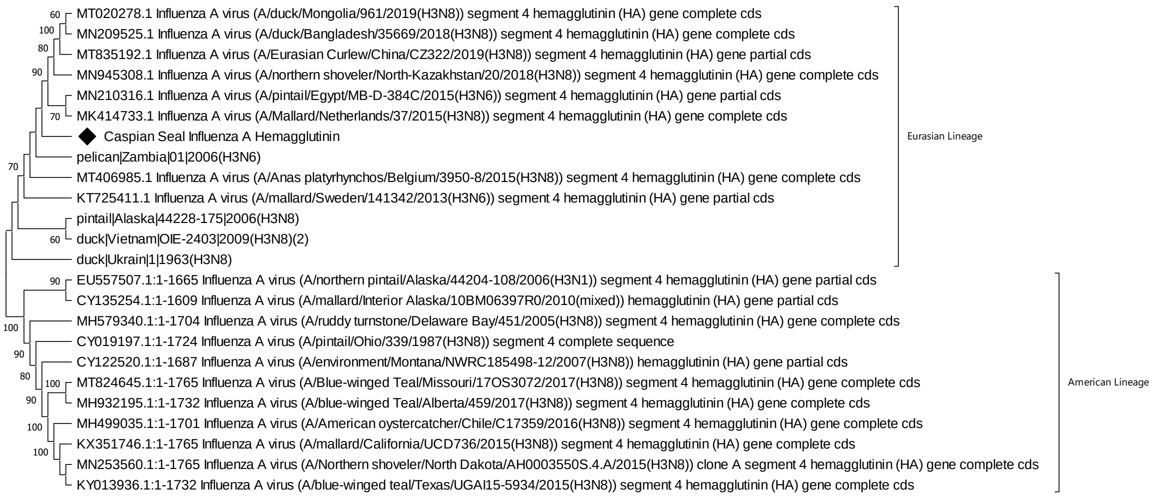
MT020278.1 Influenza A virus (A/duck/Mongolia/961/2019(H3N8)) segment 4 hemagglutinin (HA) gene complete cds
MN209525.1 Influenza A virus (A/duck/Bangladesh/35669/2018(H3N8)) segment 4 hemagglutinin (HA) gene complete cds
MT835192.1 Influenza A virus (A/Eurasian Curlew/China/CZ322/2019(H3N8)) segment 4 hemagglutinin (HA) gene partial cds
MN945308.1 Influenza A virus (A/northern shoveler/North-Kazakhstan/20/2018(H3N8)) segment 4 hemagglutinin (HA) gene complete cds
MN210316.1 Influenza A virus (A/pintail/Egypt/MB-D-384C/2015(H3N6)) segment 4 hemagglutinin (HA) gene partial cds
MK414733.1 Influenza A virus (A/Mallard/Netherlands/37/2015(H3N8)) segment 4 hemagglutinin (HA) gene complete cds
Caspian Seal Influenza A Hemagglutinin
pelican|Zambia|01|2006(H3N6)
MT406985.1 Influenza A virus (A/Anas platyrhynchos/Belgium/3950-8/2015(H3N8)) segment 4 hemagglutinin (HA) gene complete cds
KT725411.1 Influenza A virus (A/mallard/Sweden/141342/2013(H3N6)) segment 4 hemagglutinin (HA) gene partial cds
pintail|Alaska|44228-175|2006(H3N8)
duck|Vietnam|OIE-2403|2009(H3N8)(2)
duck|Ukrain|1|1963(H3N8)
Eurasian Lineage
EU557507.1:1-1665 Influenza A virus (A/northern pintail/Alaska/44204-108/2006(H3N1)) segment 4 hemagglutinin (HA) gene partial cds
CY135254.1:1-1609 Influenza A virus (A/mallard/Interior Alaska/10BM06397R0/2010(mixed)) hemagglutinin (HA) gene partial cds
MH579340.1:1-1704 Influenza A virus (A/ruddy turnstone/Delaware Bay/451/2005(H3N8)) segment 4 hemagglutinin (HA) gene complete cds
CY019197.1:1-1724 Influenza A virus (A/pintail/Ohio/339/1987(H3N8)) segment 4 complete sequence
CY122520.1:1-1687 Influenza A virus (A/environment/Montana/NWRC185498-12/2007(H3N8)) hemagglutinin (HA) gene partial cds
MT824645.1:1-1765 Influenza A virus (A/Blue-winged Teal/Missouri/17OS3072/2017(H3N8)) segment 4 hemagglutinin (HA) gene complete cds
MH932195.1:1-1732 Influenza A virus (A/blue-winged Teal/Alberta/459/2017(H3N8)) segment 4 hemagglutinin (HA) gene complete cds
MH499035.1:1-1701 Influenza A virus (A/American oystercatcher/Chile/C17359/2016(H3N8)) segment 4 hemagglutinin (HA) gene complete cds
KX351746.1:1-1765 Influenza A virus (A/mallard/California/UCD736/2015(H3N8)) segment 4 hemagglutinin (HA) gene complete cds
MN253560.1:1-1765 Influenza A virus (A/Northern shoveler/North Dakota/AH0003550S.4.A/2015(H3N8)) clone A segment 4 hemagglutinin (HA) gene complete cds
KY013936.1:1-1732 Influenza A virus (A/blue-winged teal/Texas/UGAI15-5934/2015(H3N8)) segment 4 hemagglutinin (HA) gene complete cds
American Lineage

I.

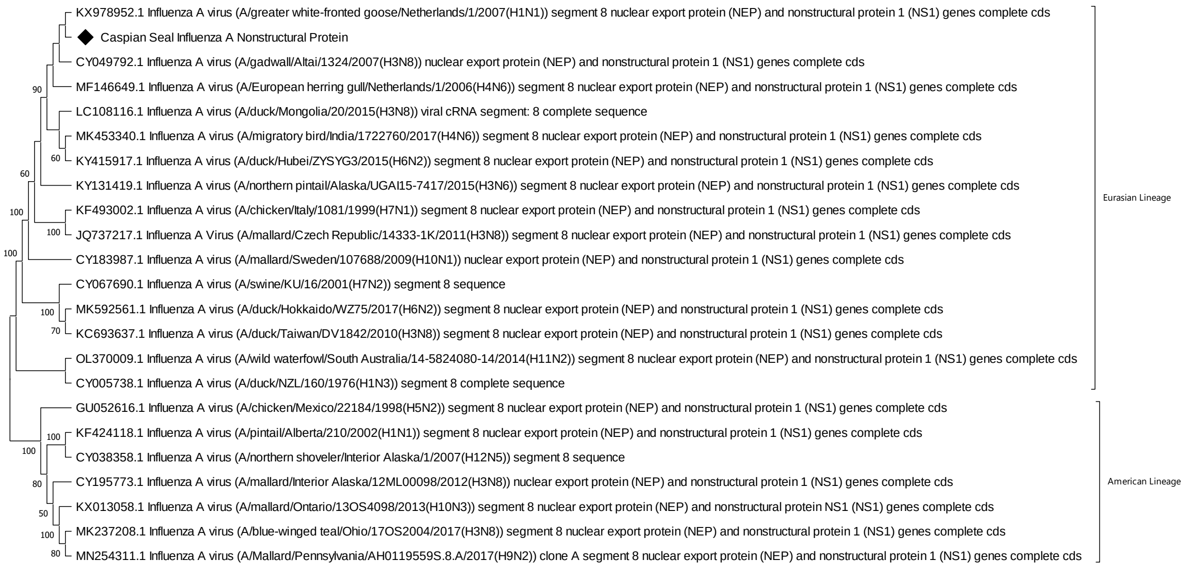
KX978952.1 Influenza A virus (A/greater white-fronted goose/Netherlands/1/2007(H1N1)) segment 8 nuclear export protein (NEP) and nonstructural protein 1 (NS1) genes complete cds
Caspian Seal Influenza A Nonstructural Protein
CY049792.1 Influenza A virus (A/gadwall/Altai/1324/2007(H3N8)) nuclear export protein (NEP) and nonstructural protein 1 (NS1) genes complete cds
MF146649.1 Influenza A virus (A/European herring gull/Netherlands/1/2006(H4N6)) segment 8 nuclear export protein (NEP) and nonstructural protein 1 (NS1) genes complete cds
LC108116.1 Influenza A virus (A/duck/Mongolia/20/2015(H3N8)) viral cRNA segment: 8 complete sequence
MK453340.1 Influenza A virus (A/migratory bird/India/1722760/2017(H4N6)) segment 8 nuclear export protein (NEP) and nonstructural protein 1 (NS1) genes complete cds
KY415917.1 Influenza A virus (A/duck/Hubei/ZYSYG3/2015(H6N2)) segment 8 nuclear export protein (NEP) and nonstructural protein 1 (NS1) genes complete cds
KY131419.1 Influenza A virus (A/northern pintail/Alaska/UGAI15-7417/2015(H3N6)) segment 8 nuclear export protein (NEP) and nonstructural protein 1 (NS1) genes complete cds
KF493002.1 Influenza A virus (A/chicken/Italy/1081/1999(H7N1)) segment 8 nuclear export protein (NEP) and nonstructural protein 1 (NS1) genes complete cds
JQ737217.1 Influenza A Virus (A/mallard/Czech Republic/14333-1K/2011(H3N8)) segment 8 nuclear export protein (NEP) and nonstructural protein 1 (NS1) genes complete cds
CY183987.1 Influenza A virus (A/mallard/Sweden/107688/2009(H10N1)) nuclear export protein (NEP) and nonstructural protein 1 (NS1) genes complete cds
CY067690.1 Influenza A virus (A/swine/KU/16/2001(H7N2)) segment 8 sequence
MK592561.1 Influenza A virus (A/duck/Hokkaido/WZ75/2017(H6N2)) segment 8 nuclear export protein (NEP) and nonstructural protein 1 (NS1) genes complete cds
KC693637.1 Influenza A virus (A/duck/Taiwan/DV1842/2010(H3N8)) segment 8 nuclear export protein (NEP) and nonstructural protein 1 (NS1) genes complete cds
OL370009.1 Influenza A virus (A/wild waterfowl/South Australia/14-5824080-14/2014(H11N2)) segment 8 nuclear export protein (NEP) and nonstructural protein 1 (NS1) genes complete cds
CY005738.1 Influenza A virus (A/duck/NZL/160/1976(H1N3)) segment 8 complete sequence
Eurasian Lineage
GU052616.1 Influenza A virus (A/chicken/Mexico/22184/1998(H5N2)) segment 8 nuclear export protein (NEP) and nonstructural protein 1 (NS1) genes complete cds
KF424118.1 Influenza A virus (A/pintail/Alberta/210/2002(H1N1)) segment 8 nuclear export protein (NEP) and nonstructural protein 1 (NS1) genes complete cds
CY038358.1 Influenza A virus (A/northern shoveler/Interior Alaska/1/2007(H12N5)) segment 8 sequence
CY195773.1 Influenza A virus (A/mallard/Interior Alaska/12ML00098/2012(H3N8)) nuclear export protein (NEP) and nonstructural protein 1 (NS1) genes complete cds
KX013058.1 Influenza A virus (A/mallard/Ontario/13OS4098/2013(H10N3)) segment 8 nuclear export protein (NEP) and nonstructural protein NS1 (NS1) genes complete cds
MK237208.1 Influenza A virus (A/blue-winged teal/Ohio/17OS2004/2017(H3N8)) segment 8 nuclear export protein (NEP) and nonstructural protein 1 (NS1) genes complete cds
MN254311.1 Influenza A virus (A/Mallard/Pennsylvania/AH0119559S.8.A/2017(H9N2)) clone A segment 8 nuclear export protein (NEP) and nonstructural protein 1 (NS1) genes complete cds
American Lineage

J.

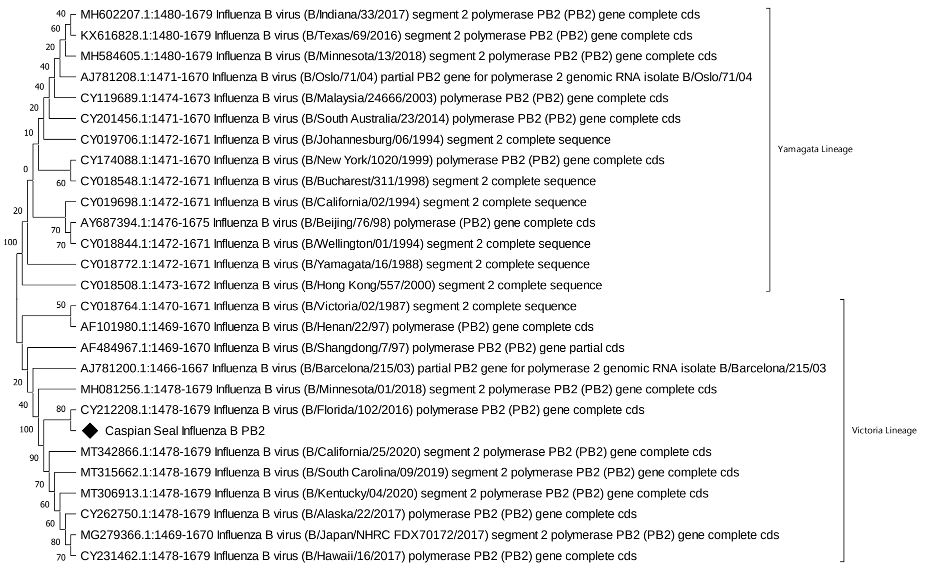
MH602207.1:1480-1679 Influenza B virus (B/Indiana/33/2017) segment 2 polymerase PB2 (PB2) gene complete cds
KX616828.1:1480-1679 Influenza B virus (B/Texas/69/2016) segment 2 polymerase PB2 (PB2) gene complete cds
MH584605.1:1480-1679 Influenza B virus (B/Minnesota/13/2018) segment 2 polymerase PB2 (PB2) gene complete cds
AJ781208.1:1471-1670 Influenza B virus (B/Oslo/71/04) partial PB2 gene for polymerase 2 genomic RNA isolate B/Oslo/71/04
CY119689.1:1474-1673 Influenza B virus (B/Malaysia/24666/2003) polymerase PB2 (PB2) gene complete cds
CY201456.1:1471-1670 Influenza B virus (B/South Australia/23/2014) polymerase PB2 (PB2) gene complete cds
CY019706.1:1472-1671 Influenza B virus (B/Johannesburg/06/1994) segment 2 complete sequence
CY174088.1:1471-1670 Influenza B virus (B/New York/1020/1999) polymerase PB2 (PB2) gene complete cds
CY018548.1:1472-1671 Influenza B virus (B/Bucharest/311/1998) segment 2 complete sequence
CY019698.1:1472-1671 Influenza B virus (B/California/02/1994) segment 2 complete sequence
AY687394.1:1476-1675 Influenza B virus (B/Beijing/76/98) polymerase (PB2) gene complete cds
CY018844.1:1472-1671 Influenza B virus (B/Wellington/01/1994) segment 2 complete sequence
CY018772.1:1472-1671 Influenza B virus (B/Yamagata/16/1988) segment 2 complete sequence
CY018508.1:1473-1672 Influenza B virus (B/Hong Kong/557/2000) segment 2 complete sequence
Yamagata Lineage
CY018764.1:1470-1671 Influenza B virus (B/Victoria/02/1987) segment 2 complete sequence
AF101980.1:1469-1670 Influenza B virus (B/Henan/22/97) polymerase (PB2) gene complete cds
AF484967.1:1469-1670 Influenza B virus (B/Shangdong/7/97) polymerase PB2 (PB2) gene partial cds
AJ781200.1:1466-1667 Influenza B virus (B/Barcelona/215/03) partial PB2 gene for polymerase 2 genomic RNA isolate B/Barcelona/215/03
MH081256.1:1478-1679 Influenza B virus (B/Minnesota/01/2018) segment 2 polymerase PB2 (PB2) gene complete cds
CY212208.1:1478-1679 Influenza B virus (B/Florida/102/2016) polymerase PB2 (PB2) gene complete cds
Caspian Seal Influenza B PB2
MT342866.1:1478-1679 Influenza B virus (B/California/25/2020) segment 2 polymerase PB2 (PB2) gene complete cds
MT315662.1:1478-1679 Influenza B virus (B/South Carolina/09/2019) segment 2 polymerase PB2 (PB2) gene complete cds
MT306913.1:1478-1679 Influenza B virus (B/Kentucky/04/2020) segment 2 polymerase PB2 (PB2) gene complete cds
CY262750.1:1478-1679 Influenza B virus (B/Alaska/22/2017) polymerase PB2 (PB2) gene complete cds
MG279366.1:1469-1670 Influenza B virus (B/Japan/NHRC FDX70172/2017) segment 2 polymerase PB2 (PB2) gene complete cds
CY231462.1:1478-1679 Influenza B virus (B/Hawaii/16/2017) polymerase PB2 (PB2) gene complete cds
Victoria Lineage

K.

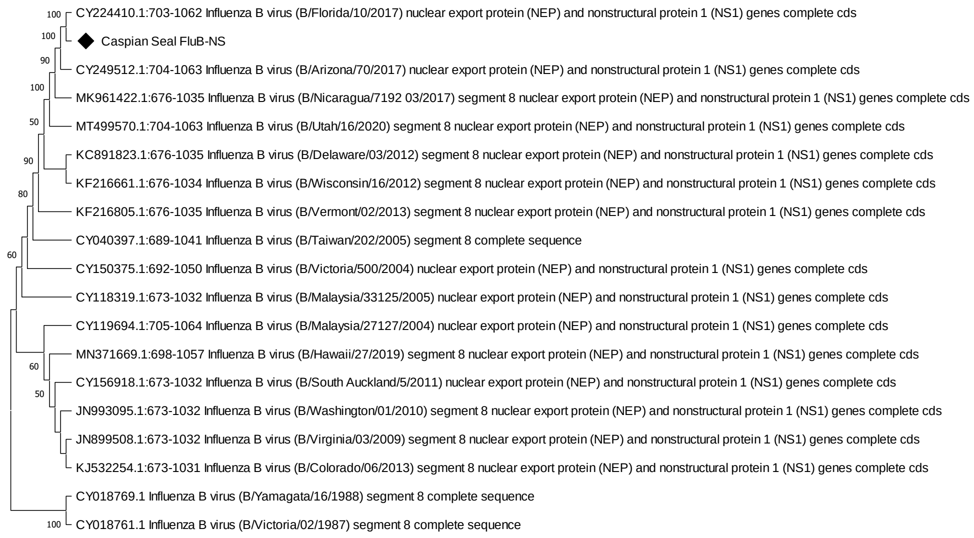


Supplementary Figure 1. Phylogenetic trees

A - Circoviridae, B – Parvoviridae, C – Herpesviridae, D – Papillomaviridae, E – Picornaviridae, F – Caliciviridae, G – Anelloviridae, H – Influenza A virus Haemagglutinin, I - Influenza A virus Non-Structural Protein, J - Influenza B virus PB2 Protein, K - Influenza B virus Non-Structural Protein
