## Supplementary TableS1 for "VIRAL METAGENOMIC SURVEY OF CASPIAN SEALS"

Supplementary Table S1. Sampling locations and dates

| Year | Month | Location | Number of seals sampled | Number of seals taken for NGS | Number of libraries constructed |
| --- | --- | --- | --- | --- | --- |
| 2009 | November | Kendirli isles, Kazakh Bay (42°44ʹ N, 52°32ʹ E) | 8 | 1 | 1 |
| 2016 | October | Kendirli isles, Kazakh Bay (42°44ʹ N, 52°32ʹ E) | 20 | 5 | 8 |
| 2019 | November | Kashagan  46°07′6" N, 52°33′8" E | 10 | 8 | 9 |
| 2020 | October | Prorva artisanal isle  46°00' N, 53°04' E | 13 | 13 | 15 |
| 2020 | December | Dagestan seashore  43°490' N, 47°462' E | 8 | 8 | 1 |
